# Are circadian amplitudes and periods correlated? A new *twist* in the story

**DOI:** 10.1101/2023.05.17.541139

**Authors:** Marta del Olmo, Christoph Schmal, Camillo Mizaikoff, Saskia Grabe, Christian Gabriel, Achim Kramer, Hanspeter Herzel

## Abstract

Three parameters are important to characterize a circadian and in general any biological clock: period, phase and amplitude. While circadian periods have been shown to correlate with entrainment phases, and clock amplitude influences the phase response of an oscillator to pulse-like zeitgeber signals, the co-modulations of amplitude and periods, which we term twist, have not been studied in detail. In this paper we define two concepts: parametric twist refers to amplitude-period correlations arising in ensembles of self-sustained clocks in the absence of external inputs, and phase space twist refers to the co-modulation of an individual clock’s amplitude and period in response to external zeitgebers. Our findings show that twist influences the interaction of oscillators with the environment, facilitating entrainment, fastening recovery to pulse-like perturbations or modifying the response of an individual clock to coupling. This theoretical framework might be applied to understand the emerging properties of other oscillating systems.

## I. Introduction

Oscillations are happening all around us, from the vibrating atoms constituting matter to the beating of the animal heart or to circadian clocks present in all kingdoms of life. Circadian clocks are autonomous clocks that tick in the absence of external timing cues with a period of about 24 h and regulate our behavior, physiology and metabolism. A fundamental property of circadian clocks is that their phase and periodicity can be adjusted to external timing signals (zeitgebers) in a process known as entrainment. It is believed that natural selection has acted on the phase relationship between biological rhythms and the environmental cycle, and thus, this phase of entrainment is of central importance for the fitness of the organism, allowing it to anticipate to changes in the external world [1].

Three key properties of a circadian rhythm are its period, amplitude and phase. Chronobiological studies have usually focused on period because there are established tools that allow their direct measurement, including running wheels for mice or race tubes for fungi. Phase, which refers to the position of a point on the oscillation cycle relative to a reference point, can be measured similarly. Circadian rhythms can be entrained to various zeitgeber periods *T* as reviewed in [2], but under the natural conditions of *T* = 24 h, variations of intrinsic periods lead to different phases of entrainment that are the basis of chronotypes: faster running clocks (shorter endogenous periods) lead to early phases (‘morning larks’) and slower clocks (longer periods) correspond to later phases (‘night owls’) [3–7].

Measuring amplitudes, however, is less straightforward. Some studies have considered activity recordings [8, 9] or conidiation in race tubes [10], but one might argue that these measures do not represent the clockwork’s amplitude, as they reflect outputs of the circadian system. Other studies have quantified gene expression profiles after careful normalization [11], but from the *∼*20 core clock genes that constitute the mammalian circadian oscillator [12], it is not immediately evident which gene or protein is best representing the core clock amplitude. Actually, reporter signals monitoring expression of different clock genes and proteins have been used to quantify amplitudes [4, 13, 14]. Others have approached the amplitude challenge indirectly by measuring the response of an oscillator to zeitgeber pulses [15–18]. While small-amplitude clocks exhibit larger pulse-induced phase shifts and are easier to phase-reset, larger amplitude rhythms display smaller phase shifts [17–22], with evident consequences in the size of the phase response curve [5] or in jet lag duration [23]. Amplitudes, together with periods, also govern entrainment [23, 24] and seasonality [22, 23, 25]. There have been various theoretical and experimental studies showing, for example, how clocks with larger amplitudes display narrower ranges of entrainment than rhythms of lower amplitude [20], and how the phase of entrainment is modulated by oscillator amplitude [22, 24].

Taken together, these observations indicate, firstly, that the phase of entrainment is correlated with the intrinsic period, and secondly, that both phase of entrainment and phase changes in response to perturbations also correlate with oscillator amplitude. This leads to the question of whether amplitudes and periods are also co-modulated and what insights these interdependencies provide about the underlying oscillator. These questions are the focus of this paper. Do faster-running clocks have larger or smaller amplitudes than slower clocks? What are the implications? Experimental observations have provided evidence for both: in a human osteosarcoma cell line in culture, clocks with longer periods display larger amplitudes [26]; but in cells from the choroid plexus, the major producer of cerebrospinal fluid of the central nervous system, clocks with shorter periods are associated with larger amplitudes [14] (scheme in Figure 1). This dependence between periods and amplitudes is what we here refer to as *twist*, also known as *shear* in the literature [27, 28]. By convention, negative twist describes oscillators in which amplitude increases are accompanied by a decreasing period (also termed hard oscillators) and *vice versa* for positive twist (soft oscillators).

**Figure 1:**
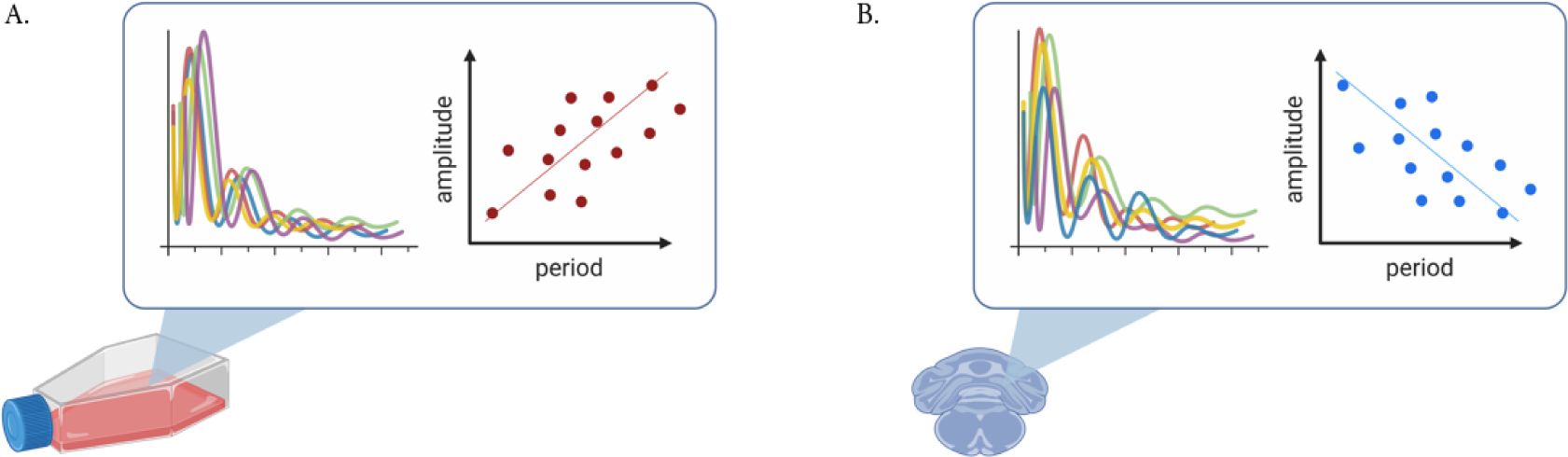
Schematic of experimental observations of circadian amplitude-period correlations. (A) Positive correlations (i.e., positive twist) have been observed in U-2 OS cells kept in culture [26]; (B) negative correlations have been observed in cells from the choroid plexus in the mouse brain [14].

In this paper we provide definitions for two important concepts: parametric twist and phase space twist. We show, firstly, that nonlinearities can introduce amplitude-period correlations in simple oscillator models. Moreover, clock models of different complexity can reproduce the experimentally-observed positive [26] and negative [14] twist effects, with the type of correlation depending on the model, on parameters, as well as on the variable being measured, illustrating the complexity in defining circadian amplitudes. Lastly, we show how twist effects can speed up or slow down zeitgeber-induced amplitude changes. This helps the clock phase adapt and modulate entrainment of oscillators to a periodic signal or their response to coupling. Our results support the use of oscillator theory as a framework to understand emerging properties of circadian clocks with different twist. Moreover, they provide insights into how temporal or spatial phase patterning might arise in coupled networks, as well as how amplitude changes could help in stabilizing the circadian period in the face of temperature changes. Although we focus on circadian clocks, the presented theory can be applied to any other oscillatory system such as cardiac rhythms, flashing fireflies or voice production.

## II. Results

### Nonlinearities can result in amplitude-period correlations across oscillators

The simple harmonic oscillator (Figure 2A) is a classical model of a system that oscillates with a restoring force proportional to its displacement, namely *F* = *−kx* [29]. The ordinary differential equation that describes the motion of a mass attached on a spring is linear (equation 1 in Materials and Methods) and the solution can be found analytically [30, 31]. The period is determined by the size of the mass *m* and the force constant *k* (see Materials and Methods), while the amplitude and phase are determined by the starting position and the velocity. Thus, an ensemble of harmonic oscillators with different initial conditions will produce results that differ in amplitudes but whose periods are the same (Figure 2B, C). When plotting amplitudes against periods, no correlation or *twist* is observed: the period of a simple harmonic oscillator is independent of its amplitude (Figure 2D).

**Figure 2:**
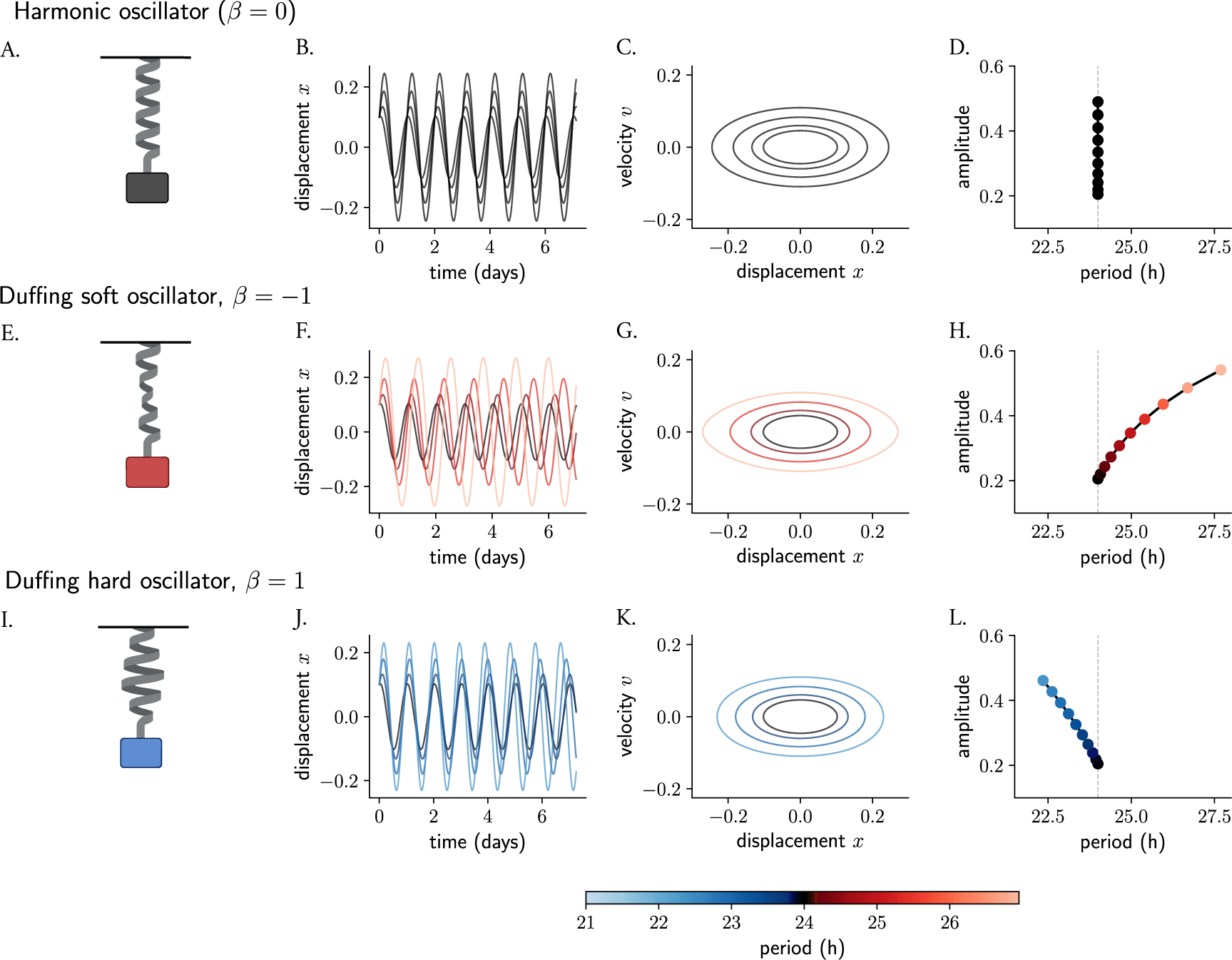
Nonlinearities introduce amplitude-period correlations in oscillator models. The mass-spring harmonic oscillator (A) is a linear oscillator that, in the absence of friction, produces persistent undamped oscillations (B, in time series and C, in phase space). Different starting conditions produce oscillations that differ in amplitudes but whose period is independent of amplitude (D). The harmonic oscillator is converted to a Duffing oscillator (E, I) by introducing nonlinearities in the restoring force of the spring (the non-linear behavior is represented with deformations of the spring). The new restoring force results in slight changes in the initial conditions producing oscillations whose amplitude and period change (F, G, J, K) and become co-dependent (H, L) and where the correlations depend on the sign of the non-linear term. The terms soft and hard oscillators refer to, by convention, those with positive and negative twist, respectively. Period values in (F, G, H, J, K, L) are color-coded.

The simple harmonic oscillator can be converted into a Duffing oscillator [32] by including a cubic nonlinearity in its equation. In the Duffing oscillator (equations 2, 3 in Materials and Methods), the restoring force is no longer linear (see deformed springs in Figure 2) but instead described by *F* = *−kx − βx*^3^, where *β* represents the coefficient of non-linear elasticity. These classical conservative oscillators instead are known to have an amplitude-dependent period. A network of Duffing clocks with different starting conditions will produce oscillations of different amplitudes as well as periods. Duffing oscillators with a negative cubic term have been termed soft oscillators and display periods that grow with amplitudes (Figure 2E-H), similar to Kepler’s Third Law of planetary motion [33], where planets with larger distances to the Sun run at slower periods than those that are closer. On the other hand, Duffing oscillators with a positive *β* term are known as hard oscillators and show negative twist (amplitude-period correlations) (Figure 2I-L) [34, 35].

In short, nonlinearities in oscillator models can introduce twist effects among ensembles of oscillators with slight differences in their properties (initial conditions, parameters, etc.). Thus, models for the circadian clock, which are based on nonlinearities, are expected to show amplitude-period correlations.

### Oscillator heterogeneity produces parametric twist effects in limit cycle clock models

Most circadian clock models generate stable limit cycle oscillations. Limit cycles are isolated closed periodic orbits with a given amplitude and period, where neighboring trajectories (e.g. any perturbation applied to the cycle) spiral either towards or out of the limit cycle [36, 37]. Stable limit cycles are examples of attractors: they imply self-sustained oscillations. The closed trajectory describes the perfect periodic behavior of the system, and any small perturbation from this trajectory causes the system to return to it, to be *attracted back* to it.

Limit cycles are inherently non-linear phenomena and they cannot occur in a linear system (i.e., a system in the form of 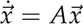, like the harmonic oscillator). From the previous section we have learned that non-linear terms can introduce amplitude-period co-dependencies. In this section we show that kinetic limit cycle models of the circadian clock also show twist effects. Instead of studying the amplitude-period correlation of an oscillator model with fixed parameters and changing initial conditions as in Figure 2 (as all initial conditions would be attracted to the same limit cycle), we study here the correlations that arise among *different* uncoupled oscillators due to oscillator heterogeneity (i.e., differences in biochemical parameters), as found experimentally [14, 26]. We refer to amplitude-period correlations that arise due to oscillator heterogeneity as *parametric twist*.

#### Single negative feedback loop models

The Goodwin model is a simple kinetic oscillator model [38] that is based on a delayed negative feedback loop, where the final product of a 3-step chain of reactions inhibits the production of the first component (equation 4 in Materials and Methods). In the context of circadian rhythms [39, 40], the model can be interpreted as a clock activator *x* that produces a clock protein *y* that, in turn, activates a transcriptional inhibitor *z* that represses *x* (Figure 3A). The Goodwin model has been extensively studied and fine-tuned by Gonze and others [41] to study fundamental properties of circadian clocks [42–45] or synchronization and entrainment [41, 46, 47]. The Gonze model [41] (equation 5) includes additional nonlinearities, where the degradation of all 3 variables is modeled with non-linear Michaelis Menten kinetics, to reduce the need of very large Hill exponents (*n* > 8), required in the original Goodwin model to generate self-sustained oscillations [48], that have been questioned to be biologically meaningful. These Michaelian degradation processes can be interpreted as positive feedback loops which aid in the generation of oscillations [49] (Figure 3B).

**Figure 3:**
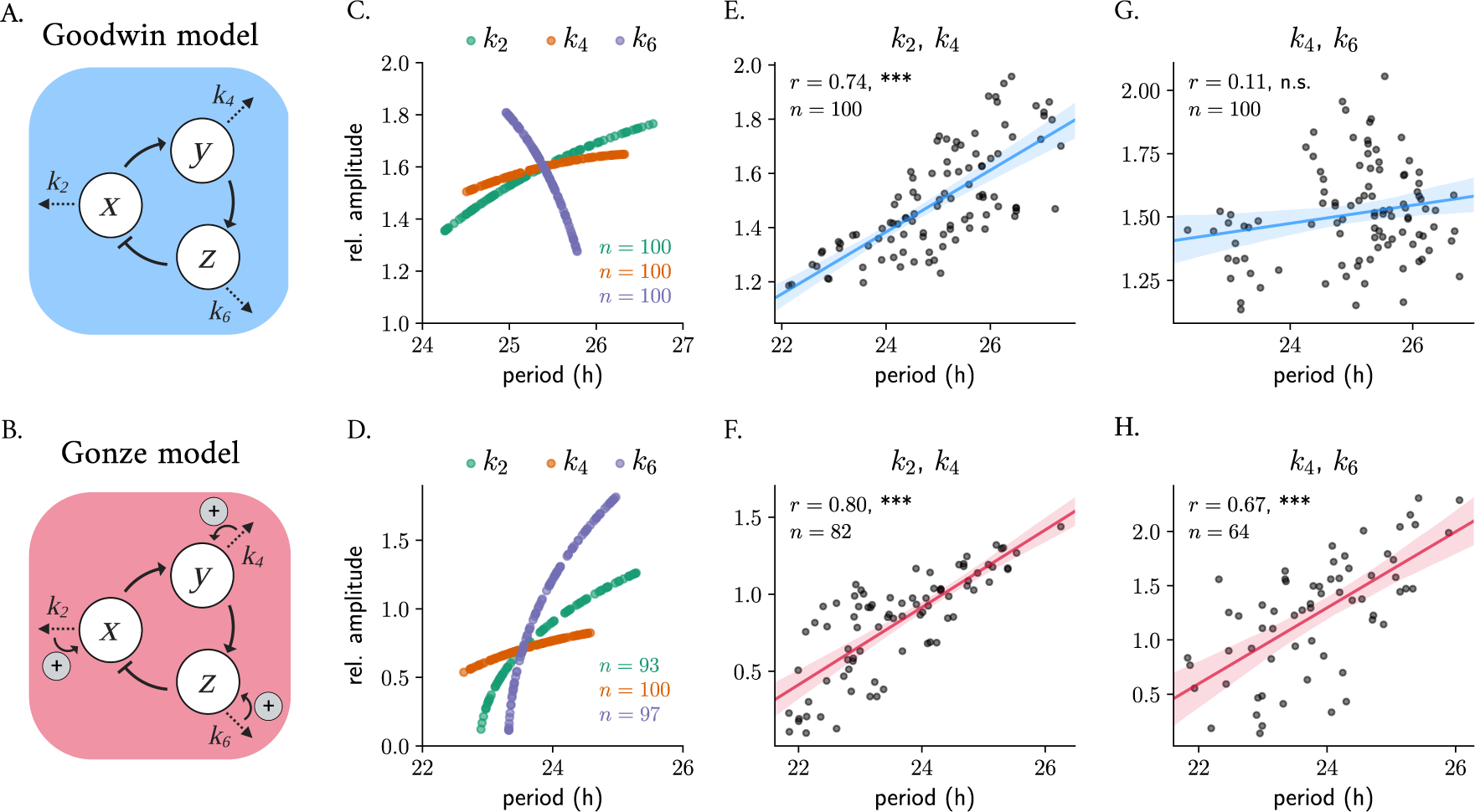
Parametric twist in an ensemble of 100 uncoupled Goodwin-like models: correlations are model- and parameter-dependent. Scheme of the Goodwin (A) and Gonze (B) oscillator models. Changes in degradation parameters result in twist effects when varied randomly around *±*10% their default parameter value, mimicking oscillator heterogeneity of the Goodwin (C) and Gonze (D) models. Simultaneous co-variation of degradation parameters results in parametric twist: changes of *k*_2_ and *k*_4_ produce significant positive amplitude-period correlations in both models (E, F); co-variations of *k*_6_ and *k*_4_ result in amplitude-period correlations that are not significant in the Goodwin model (G), but significant positive parametric twist in the Gonze model (H). Shown in all panels is Spearman’s *R* and the significance of the correlation (n. s.: not significant; *^∗∗∗^*: *p* value<0.001). Default model parameters are summarized in Table 1. Relative amplitudes are computed as the mean peak-to-trough distance of the *x* variable. Oscillators from the ensemble whose amplitude was < 0.1 have been removed from the plots; the total number of oscillators (*n*) is indicated in the plots.

**Table 1:**
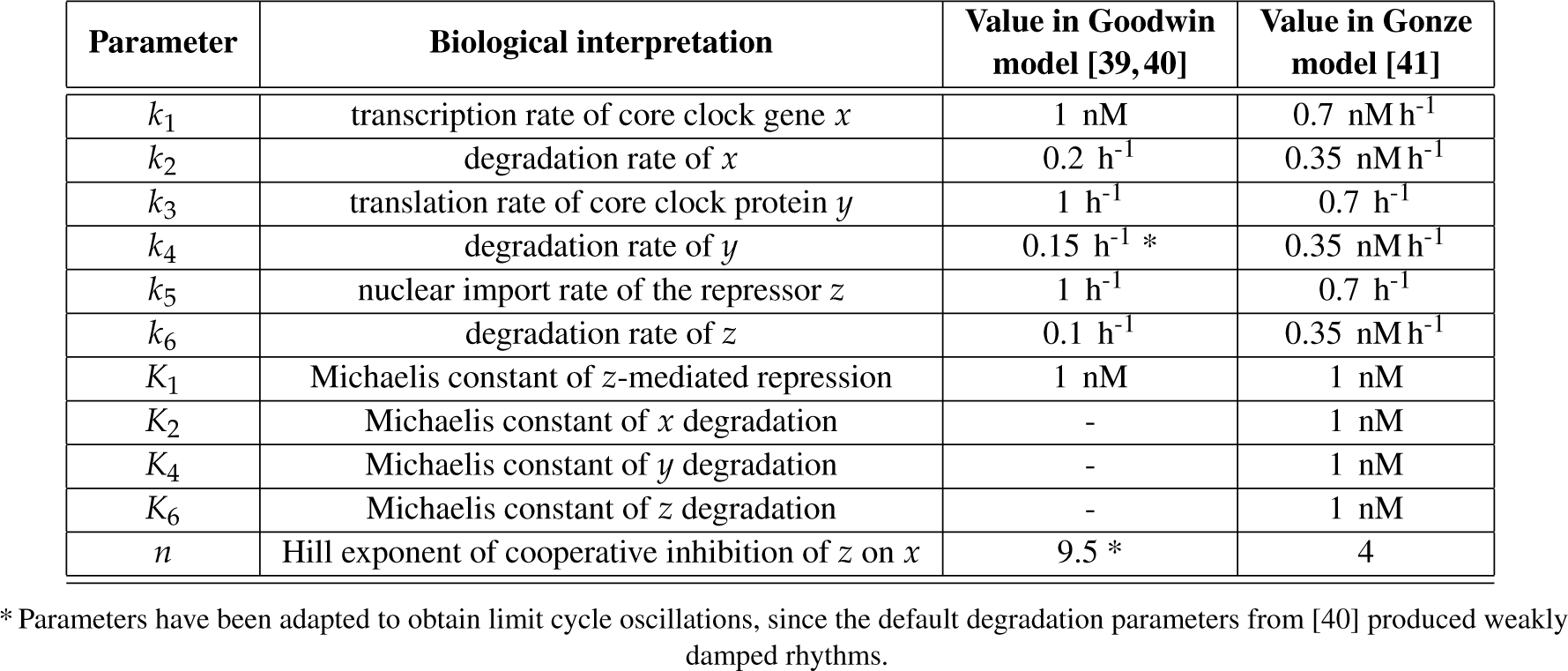
Default parameters of the Goodwin and Gonze models used for our simulations. Their biological interpretation as well as the default values from publications are shown.

| Parameter | Biological interpretation | Value in Goodwin model [39, 40] | Value in Gonze model [41] |
| --- | --- | --- | --- |
| $k_1$ | transcription rate of core clock gene $x$ | 1 nM | 0.7 nM h<sup>-1</sup> |
| $k_2$ | degradation rate of $x$ | 0.2 h<sup>-1</sup> | 0.35 nM h<sup>-1</sup> |
| $k_3$ | translation rate of core clock protein $y$ | 1 h<sup>-1</sup> | 0.7 h<sup>-1</sup> |
| $k_4$ | degradation rate of $y$ | 0.15 h<sup>-1</sup> * | 0.35 nM h<sup>-1</sup> |
| $k_5$ | nuclear import rate of the repressor $z$ | 1 h<sup>-1</sup> | 0.7 h<sup>-1</sup> |
| $k_6$ | degradation rate of $z$ | 0.1 h<sup>-1</sup> | 0.35 nM h<sup>-1</sup> |
| $K_1$ | Michaelis constant of $z$-mediated repression | 1 nM | 1 nM |
| $K_2$ | Michaelis constant of $x$ degradation | - | 1 nM |
| $K_4$ | Michaelis constant of $y$ degradation | - | 1 nM |
| $K_6$ | Michaelis constant of $z$ degradation | - | 1 nM |
| $n$ | Hill exponent of cooperative inhibition of $z$ on $x$ | 9.5 * | 4 |

\* Parameters have been adapted to obtain limit cycle oscillations, since the default degradation parameters from [40] produced weakly damped rhythms.

To study whether twist effects are present in an ensemble of 100 uncoupled Goodwin and Gonze oscillators with different parameter values, we randomly varied the degradation parameters of *x*, *y* or *z* (*k*_2_, *k*_4_ and *k*_6_, respectively) in each oscillator around *±*10% their default parameter value (given in Table 1). The resulting parametric twist depends on the model and on the parameter being changed: in the Goodwin model, variations in the degradation rate of the transcriptional activator (*k*_2_) or of the clock protein (*k*_4_) result in positive twist effects (i.e., soft twist-control), while changes in the transcriptional repressor’s degradation rate (*k*_6_) result in a hard twist-control, i.e., negative amplitude-period correlation (Figure 3C). In the Gonze model, *k*_2_, *k*_4_ and *k*_6_ are all soft twist-control parameters, as individual changes of any of them all produce positive parametric twist effects (Figure 3D).

To mimic cell-to-cell variability in a more realistic manner, we introduced heterogeneity by changing combinations of the degradation parameters simultaneously around *±*10% their default parameter value and analyzing the resulting periods and amplitudes. We observed that the overall twist behavior depends on the particular influence that each parameter has individually. Random co-variations of *k*_2_ and *k*_4_ produce significant positive parametric twist effects in ensembles of both Goodwin (Figure 3E) or Gonze clocks (Figure 3F), consistent with the positive correlations when either of the parameters is changed individually (Figure 3C, D). Nevertheless, when *k*_6_, which has a negative twist effect in the Goodwin model, is changed at the same time as *k*_4_ (or *k*_2_, data not shown) the correlation is not significant (Figure 3G). Random co-variation of *k*_4_ and *k*_6_ in an ensemble of Gonze clocks results in significant positive parametric twist effects (Figure 3H).

Changes in parameter values can result in significant alterations to a system’s long-term behavior, which can include differences in the number of steady-states, limit cycles, or their stability properties. Such qualitative changes in non-linear dynamics are known as bifurcations, with the corresponding parameter values at which they occur being referred to as bifurcation points. In oscillatory systems, Hopf bifurcations are an important type of bifurcation point. They occur when a limit cycle arises from a stable steady-state that loses its stability. A 1-dimensional bifurcation diagram illustrates how changes in a mathematical model’s control parameter affect its final states, for example the period or amplitude of oscillations. The Hopf bubble refers to the region in parameter space where the limit cycle exists, and it is commonly represented with the peaks and troughs of a measured variable in the *y* axis, with the control parameter plotted on the *x* axis (Figure 4A). The term “bubble” is used because of the shape of the curve, that resembles a bubble that grows or shrinks as the parameter is changed. Such bifurcation analyses can be used to predict the type of twist that a system shows: the amplitude will increase with parameter changes if the default parameter value is close to the opening of the Hopf bifurcation, or will decrease if the value is near the closing of the bubble.

**Figure 4:**
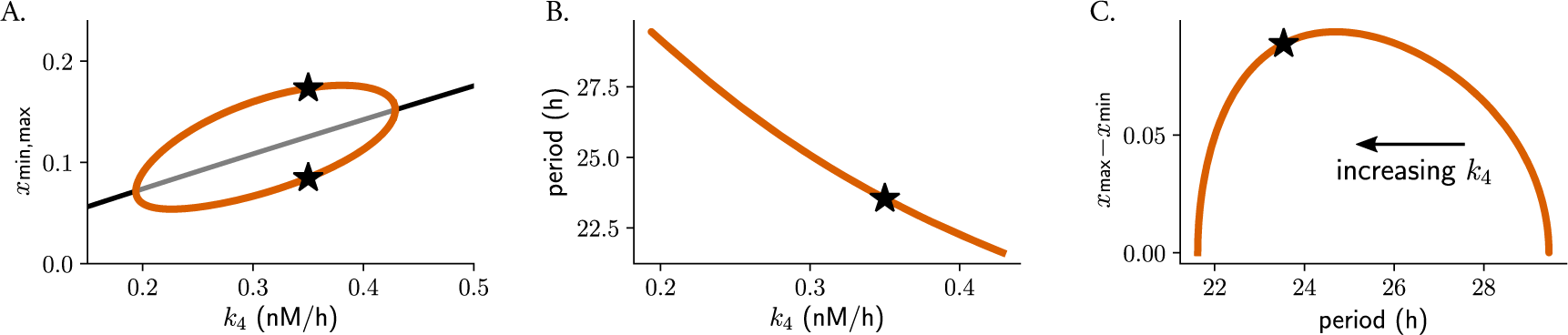
The type of parametric twist depends on the parameter location within the Hopf bubble. Bifurcation analysis of Gonze oscillators with changing *k*_4_, simulating heterogeneous oscillators: (A) Hopf bubble representing the maximum and the minimum of variable *x* as a function of changes in the degradation rate of *y* (*k*_4_), (B) monotonic period decrease for increasing *k*_4_. Represented in (C) is the combination of (A) and (B), namely how the difference between the maxima and minima of *x* changes with the period: a non-monotonic behavior of parametric twist is observed, which depends on the values of *k*_4_. Stars indicate the default parameter value, period or amplitude. Note that, since *k*_4_ results in a decrease in period length, the *k*_4_ dimension is included implicitly in (C) and *k*_4_ increases from the right to the left of the plot.

We observed that Gonze oscillators display self-sustained oscillations for values of *k*_4_ between 0.2 and 0.43 (Figure 4A) and that, for increasing *k*_4_, periods decrease monotonically (Figure 4B). For oscillators with increasing *k*_4_, the parametric twist effects from the ensemble are first negative (amplitudes increase and periods decrease); however, when Gonze clocks have *k*_4_ values that are part of the region of the bubble where amplitudes decrease, positive amplitude-period correlations appear in the ensemble (Figure 4C). In summary, the type of parametric twist depends on both the model and the parameter being studied but also on where the parameter is located within the Hopf bubble. Other more complex kinetic models of the mammalian circadian clockwork have shown that changes in some of the degradation parameters produce non-monotonic period changes [50], and thus the twist picture is expected to become even more complex.

#### Model with synergistic feedback loops

The Almeida model [51] is a more detailed model of the core clock network in mammals that includes seven core clock proteins that exert their regulation at E-boxes, D-boxes and ROR binding elements (RORE) through multiple positive and negative feedback loops (Figure 5A, equation 6 in Materials and Methods). To study the parametric twist in an ensemble of uncoupled Almeida oscillators, we randomly changed all 18 parameters individually around *±*20% their default value (Table 2 in Materials and Methods) and computed the amplitude-period correlations (i.e., parametric twist). We found, interestingly, that the overall twist effects depend not only on the parameter being studied, but also on the variable which is measured. For example, changes in the rate of D-box activation parameter *V_D_* result in negative twist effects for BMAL1 (i.e., lower amplitude BMAL1 rhythms run slower than oscillators with higher amplitude BMAL1 rhythms), positive parametric twist for the PER:CRY complex but almost no parametric twist from the perspective of PER (Figure 5B-D). Changes in PER degradation *γ_P_* produce positive parametric twist for BMAL1, PER and PER:CRY but of different magnitudes (Figure 5E-G). Supplementary Table S1 summarizes the parametric twist effects for the additional parameters from the Almeida model, highlighting the complexity that arises with synergies of feedback loops and bringing us again to the question of defining what the relevant amplitude of a complex oscillator is.

**Figure 5:**
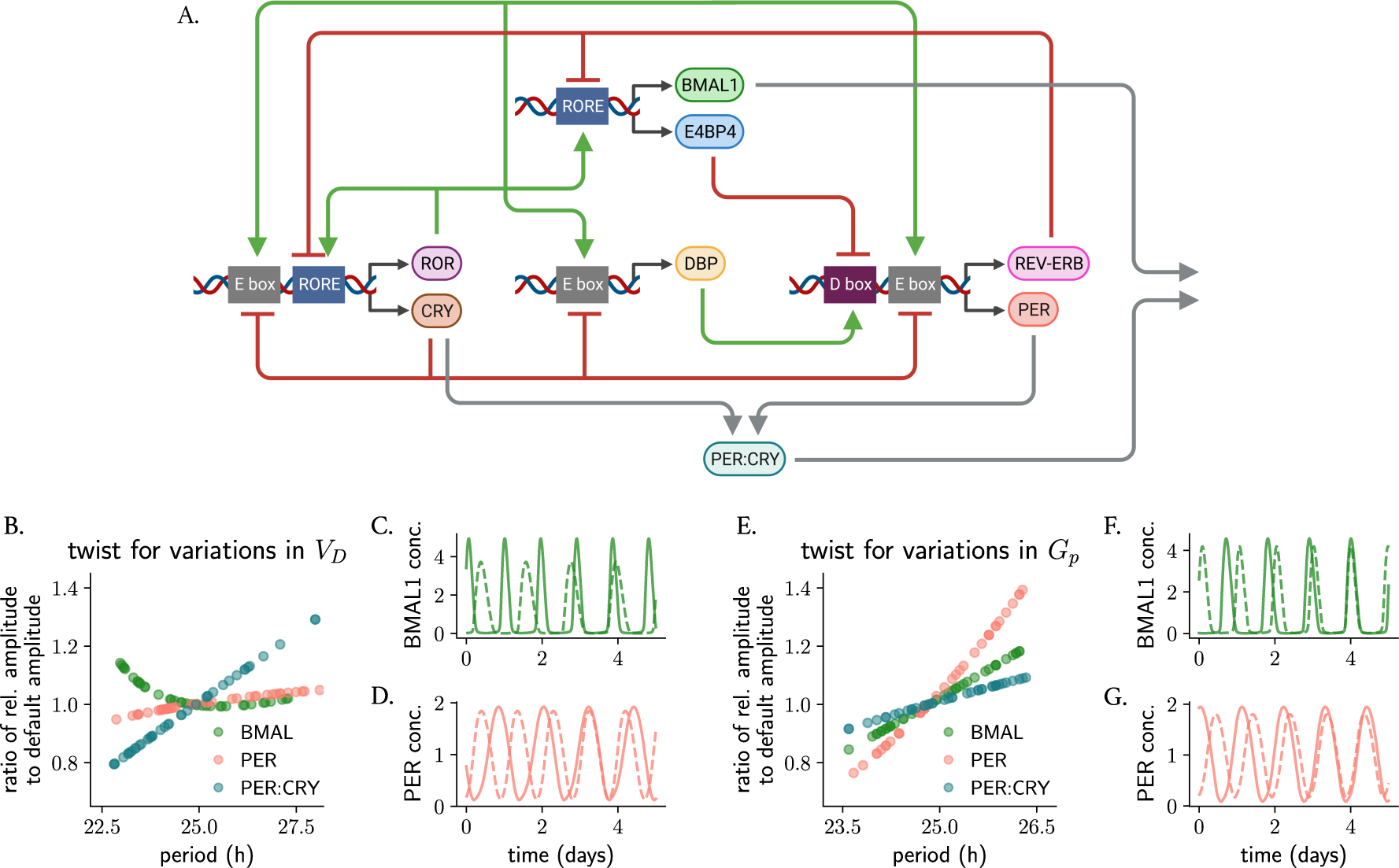
The type of parametric twist in models of the circadian clockwork with synergistic feedback loops depends on the parameter being studied and on the variable that is measured. (A) Scheme of the Almeida model [51], illustrating activation and inhibition of different clock-controlled elements (at E-boxes, D-boxes and ROR elements) by the respective core clock proteins. Parametric twist was studied by modeling a heterogeneous uncoupled ensemble where model parameters were randomly and individually varied around *±*20% of their default value (shown in Table 2) and assessing the period-amplitude correlation of the ensemble. (B-D) Almeida oscillators (*n* = 40) with heterogeneous values in the activation rate of D-boxes (*V_D_*): the twist is negative from the perspective of BMAL1, positive from the perspective of PER:CRY, but the amplitude of PER is not greatly affected by changing *V_D_*. (E-G) Almeida oscillators (*n* = 40) with heterogeneous values in the degradation rate of PER (*γ_P_*) show positive parametric twist effects for BMAL1, PER and PER:CRY. The twist in panels (B, E) is evaluated by comparing the relative amplitude of the respective protein/protein complex (computed as the average peak-to-trough distance) after parameter change to the default amplitude. Shown in (C, D, F, G) are representative oscillations with long and short periods: dashed lines represent rhythms of smallest amplitudes.

**Table 2:** List of parameters of the Almeida model. Time units are given in hours and concentration units as arbitrary units. The default values are taken from [51].

| Parameter | Biological interpretation | Value |
| --- | --- | --- |
| $V_R$ | rate of RORE activation | 44.4 h<sup>-1</sup> |
| $k_R$ | strength of RORE activation | 3.54 |
| $k_{Rr}$ | strength of RORE inhibition | 80.1 |
| $V_E$ | rate of E box activation | 30.3 h<sup>-1</sup> |
| $k_E$ | strength of E box activation | 214 |
| $k_{Er}$ | strength of E box inhibition | 1.24 |
| $V_D$ | rate of D box activation | 202 h<sup>-1</sup> |
| $k_D$ | strength of D box activation | 5.32 |
| $k_{Dr}$ | strength of D box inhibition | 94.7 |
| $\gamma_{Ror}$ | ROR degradation rate | 2.55 h<sup>-1</sup> |
| $\gamma_{Rev}$ | REV degradation rate | 0.241 h<sup>-1</sup> |
| $\gamma_P$ | PER degradation rate | 0.844 h<sup>-1</sup> |
| $\gamma_C$ | CRY degradation rate | 2.34 h<sup>-1</sup> |
| $\gamma_{DB}$ | DBP degradation rate | 0.156 h<sup>-1</sup> |
| $\gamma_{E4}$ | E4BP4 degradation rate | 0.295 h<sup>-1</sup> |
| $\gamma_{PC}$ | PER:CRY degradation rate | 0.191 h<sup>-1</sup> |
| $\gamma_{CP}$ | PER:CRY formation rate | 0.141 h<sup>-1</sup> |
| $\gamma_{BP}$ | BMAL nuclear export rate | 2.58 h<sup>-1</sup> |

### Phase space twist in single oscillators: On amplitude-phase models, isochrones and perturbed trajectories

Up until now, our results have focused on studying amplitude-correlations among ensembles of self-sustained oscillators with differences in their intrinsic properties (e.g. biochemical parameters) in the absence of external cues. However, when a clock is exposed to external stimuli, its amplitude and period undergo adaptation. We refer to these amplitude-period correlations in individual clocks as they return to their steady-state oscillation after a stimulus as *phase space twist*. In contrast to *parametric twist*, which emerged as a result of a *heterogenous population* of oscillators, here we focus on *individual* clocks.

To further explore the correlation and interdependence between the frequency of an oscillator and its amplitude upon an external stimulus, more generalized models can be of use. The Poincaré oscillator model (equation 7 in Materials and Methods) is a simple conceptual oscillator model with only two variables, amplitude and phase, that has been widely used in chronobiology research since the early 80s [20, 22, 24, 37, 52, 53]. This amplitude-phase model, regardless of molecular details, can capture the dynamics of an oscillating system and what happens when perturbations push the system away from the limit cycle. When a pulse is applied and an oscillating system is ‘kicked out’ from the limit cycle, the perturbed trajectory is attracted back to it at a rate *λ*. Within the limit cycle, the dynamics are strictly periodic: if one takes a point in phase space and observes where the system returns to after exactly one period, the answer is trivial: to exactly the same spot (Supplementary Figure S1, red dots). Nevertheless, outside the limit cycle, during the transient relaxation time (i.e., time between the perturbation and the moment that the trajectory reaches the limit cycle), the time between two consecutive peaks might be shorter or longer than the period of the limit cycle orbit. We will come back to this point in the next section.

Winfree introduced a practical term, isochrones, to conceptualize timing relations in oscillators perturbed off their attracting cycles [52], which becomes important in physiological applications because often, biological oscillators are not on their attracting limit cycles. To understand what isochrones are, Winfree proposes in [52] a simple experiment where a pulse-like perturbation applied in a system produces a ‘bump’ in the limit cycle. As the perturbation relaxes and spirals towards the attracting limit cycle, one records the position of the oscillator in time steps which equal the period of the unperturbed limit cycle. The sequence of points left as a footprint will converge to a fixed point on the cycle and will outline an isochrone (for a detailed explanation, see Supplementary Figure S1). Isochrones are directly related to oscillator twist, and whereas in an oscillator with no twist, the isochrones are straight and radial (Supplementary Figure S1A), when twist is present, the isochrones become bent or skewed (Supplementary Figure S1B, C).

In an individual oscillator with positive twist, a perturbation that increases the oscillator’s instantaneous amplitude (Supplementary Figure S1B) will relax back and intersect the isochrone at an angle > 360*^◦^* than an oscillator with no twist in the time of one period, whereas an oscillator with negative twist will cover an angle < 360*^◦^* in the time of one period (Supplementary Figure S1C). Thus, isochrones and consequently twist illustrate how a perturbation from the limit cycle of the system accelerates or decelerates the oscillation during the course of relaxation.

Mathematically, the ‘skewness’ in isochrones can be represented in an amplitude-phase mathematical model with a twist parameter ε that adds a radius dependency on the phase dynamics. The phase changes at a constant rate 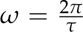 in the limit cycle, but in conditions outside the limit cycle, the phase is made to be modulated by the radius through the twist parameter *ε* until *r* = *A* (equation 7 in Materials and Methods). This *phase space twist* parameter *ε* now represents the amplitude-period correlations (acceleration/deceleration) as an *individual* oscillator returns to its steady-state oscillations.

### Phase space twist in single oscillators affects their interaction with the environment

We have seen how, in an individual oscillator, the twist parameter *ε* acts to speed up (in case of positive twist) or to slow down (in case of negative twist) trajectories further away from the limit cycle compared to those closer to it. As a result, *ε* has a direct consequence on how the individual oscillator responds to pulse-like perturbations or periodic zeitgebers coming from the environment.

To analyze how phase space twist affects the response of an oscillator to a zeitgeber pulse, we applied a pulse-like perturbation to individual Poincaré oscillators with different phase space twist values *ε*. The pulse was applied at CT3 and was made to increase the instantaneous amplitude (*A > r*). If the oscillator has no twist (*ε* = 0, Figure 6A), the isochrones are straight and radial, but for a soft or a hard oscillator with positive or negative *ε* values, respectively, isochrones get skewed (Figure 6B, C, also Supplementary Figure S1). In the case of a soft oscillator with positive *ε* (Figure 6B), the pulse at CT3 gets accelerated during the course of its relaxation, and as a consequence, the oscillator arrives back to the limit cycle at an earlier phase than the oscillator with *ε* = 0 (Figure 6D, E). The same can also be inferred mathematically: for this particular perturbation where *A > r*, *A − r >* 0 holds and hence the phase velocity outside the limit cycle is larger (*φ > ω*) for the oscillator with positive twist, since *φ* = *ω* + *ε*(*A − r*) (equation 7). The opposite holds for the oscillator with negative twist: this clock arrives at a later phase to the limit cycle (Figure 6C, F).

**Figure 6:**
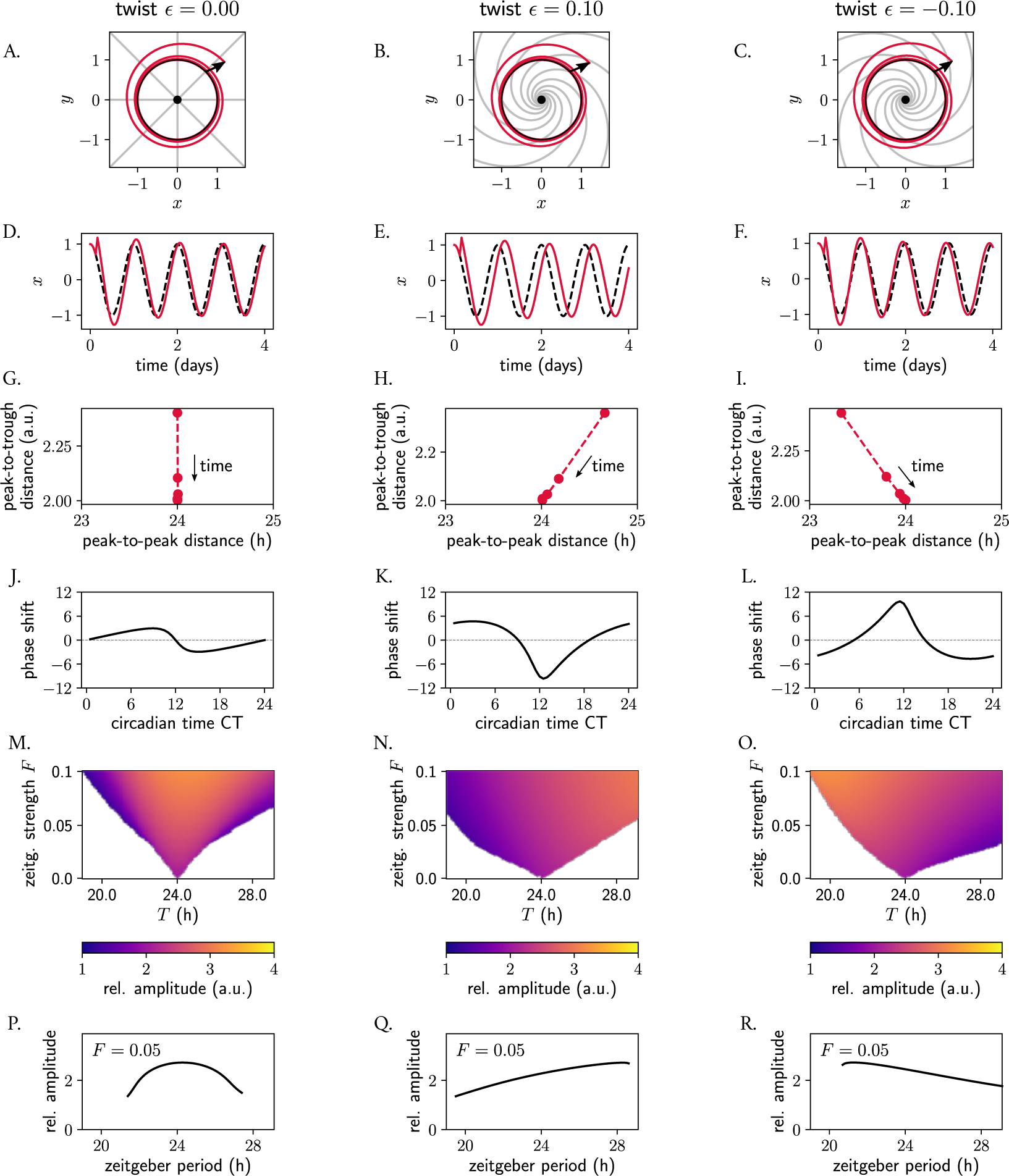
Phase space twist in a single oscillator and its effect on responses to pulse-like perturbations and to entrainment to periodic zeitgebers. (A-C) Poincaré oscillator models for different twist values *ε*, shown in phase space. The limit cycle is shown in black; perturbed trajectories are shown in red; isochrones are depicted in grey. (D-F) Time series corresponding to panels (A-C): the unperturbed limit cycle oscillations are shown with a dashed black line; perturbed trajectories are shown in red. (G-I) Peak-to-trough distance as a function of the peak-to-peak distance from the perturbed time series from (D-F) relaxing back to the limit cycle (the implicit time dimension is indicated in the panels). (J-L) Phase response curves: the shape of the PRC and the extent of the phase shifts depends on the twist *ε*. (M-O) Arnold tongues illustrating entrainment ranges of an individual oscillator with different phase space twist values to a periodic sinusoidal zeitgeber input of different periods *T*. The amplitude of the entrained clock is color-coded, with yellow colors corresponding to larger amplitudes. (P-R) The amplitude of an individual oscillator driven by a sinusoidal zeitgeber also depends on the twist of the individual oscillator. Shown are the resonance curves as a response to a zeitgeber input of strength *F* = 0.05 and varying periods *T*. Amplitudes are defined as the average peak-to-trough distances of the entrained signal.

The positive and negative amplitude-period correlations characteristic of twist effects are evident in how, upon the pulse-like perturbation, the peak-to-peak distance changes (compared to the limit cycle period) as the perturbation returns to the periodic orbit. For a clock with *ε* = 0, the peak-to-peak distance outside the limit cycle after a perturbation coincides with the 24 h peak-to-peak distance within the periodic orbit (Figure 6G). In the case of the Poincaré oscillator with positive twist, the peak-to-peak distance of the oscillator after the perturbation that increases the instantaneous amplitude is *shorter* than the period of the steady-state rhythm (24 h). As the perturbed amplitude decreases and is attracted back to the stable cycle, the peak-to-peak distance approaches 24 hours (Figure 6H, consistent with the formulation ‘positive’ twist, which implies positive amplitude-period correlations). Conversely, the clock with negative twist exhibits a longer peak-to-peak distance than the 24 h rhythm during the relaxation time, which subsequently decreases as the system returns to the stable periodic orbit (Figure 6I). When considering the findings from the last two paragraphs as a whole, it becomes clear how the extent of the phase shift between a perturbed and an unperturbed oscillator (red and black dashed lines in Figure 6D-F, respectively) depends on *ε* and thus the shape of the phase response curve (PRC) and magnitude of the phase shift depend on this twist parameter (Figure 6J-L).

We then evaluated how twist affects the response of an individual oscillator to a periodic zeitgeber input. We observed how the range of entrainment becomes larger with larger values of absolute value *ε*, as seen by the wider Arnold tongues in Figure 6M-O. It is widely known that, when the frequency of an applied periodic force is equal or close to the natural frequency of the system on which it acts, the amplitude of the oscillator on which the driving force (zeitgeber) acts on increases due to resonance effects. Interestingly, also the resonance curve was affected by twist: for the oscillator with no twist, the maximum amplitude occurred as expected for a zeitgeber period of 24 h, matching the intrinsic oscillator’s frequency (Figure 6P). In oscillators with positive or negative twist, however, the maximum amplitude after zeitgeber forcing increases and decreases with zeitgeber period, respectively (Figure 6Q, R), resulting in skewed resonance curves [54]. Interestingly, we observed that twist also affects the entrainment of an individual clock, and oscillators with high absolute values of twist cannot entrain to a periodic signal (Supplementary Figure S2), a phenomenon that has previously been described as shear-induced chaos [27, 55, 56] and that arises because the isochrones become so skewed, that the trajectory of the relaxation gets stretched and folded.

### Relaxation rate affects the response of oscillators to perturbations

Not only the twist parameter *ε*, but also the amplitude relaxation rate *λ* affects the response of oscillators to perturbations and consequently the entrainment range. We have seen how, in the Poincaré model (equation 7), the twist parameter *ε* dictates how much trajectories are ‘sped up’ or ‘slowed down’ dependent on their distance from the limit cycle. The amplitude relaxation rate *λ* describes the rate of attraction back to the limit cycle, which is independent of *ε*. Together, these parameters both dictate how ‘skewed’ the isochrones are in phase space (see analytical expression in Materials and Methods, equation 11). For Poincaré oscillators with twist, the isochrones become more radial and straight as the amplitude relaxation rate increases (Supplementary Figure S3A-C), with the implications that this has on PRCs and entrainment that have been mentioned before. A more ‘plastic’ clock (with a lower *λ* value) responds to an external pulse with larger phase shifts than a more rigid clock (with higher *λ*), resulting in phase response curves of larger amplitude (Supplementary Figure S3D-F). This is also intuitive, as it is clear that for larger values of *λ*, the twist has shorter effective time to act upon perturbed trajectories because the perturbation spends less time outside the limit cycle.

### Single oscillator phase space twist affects coupled networks

Biological oscillators are rarely alone and uncoupled. Coupled oscillators, instead, are at the heart of a wide spectrum of living things: pacemaker cells in the heart [37], insulin- and glucagon-secreting cells in the pancreas [57] or neural networks in the brain and spinal cord that control rhythmic behavior as breathing, running and chewing [58]. A number of studies have shown that networks of coupled oscillators behave in a fundamentally different way than ‘plain’ uncoupled oscillators [41, 59, 60]. This suggests that single oscillator twist might also affect how coupling synchronizes a network of oscillators.

To analyze the effect of twist on coupled oscillators, we study systems of Poincaré oscillators coupled through a mean-field (equation 8). We start by coupling two *identical* oscillators and analyzing the effect that different twist values have on the behavior of the coupled network. The numerical simulations show that increasing coupling strengths affect the period of the coupled system: oscillators with positive twist show longer periods as the coupling strength increases, whereas oscillators with negative twist show period shortening (Figure 7A and Supplementary Figure S4A-C). Coupling results in an increase in amplitude but, for our default relaxation rate value (*λ* = 0.05 h^-1^), no significant differences across oscillators with different twist values were found (Supplementary Figure S4). Increasing relaxation rates (oscillators that are attracted faster back to the limit cycle upon a perturbation) nevertheless resulted in less coupling-induced period or amplitude changes (Supplementary Figure S5).

**Figure 7:**
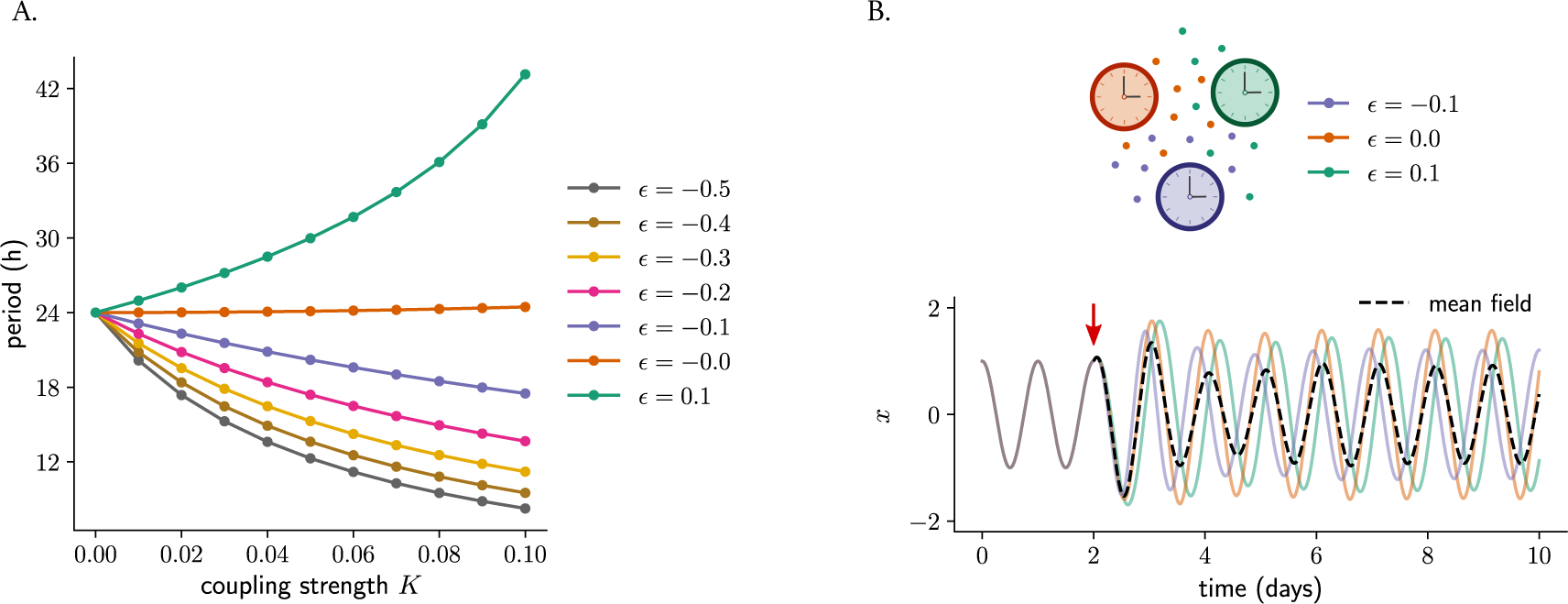
Phase space twist *ε* affects the response of a network to coupling. (A) Mean-field coupling of two identical Poincaré oscillators (sharing the same value of twist *ε*) and the effect on the period of the coupled system: whereas coupling among oscillators with positive twist results in longer periods of the coupled network, mean-field coupling among oscillators with negative twist results in the network running at a faster pace (period shortening). (B) Mean-field coupling can synchronize a network of oscillators with different twist values. Shown are three oscillators that only differ in their twist parameters (but else identical: amplitude, amplitude relaxation rate and period are the same for the three oscillators). It is seen how all oscillations overlap during the first part of the time series, since twist has no effect within the limit cycle. At a certain point (indicated by the red arrow), mean-field coupling is turned on and it is observed how the oscillators respond differently to that mean-field coupling, ending up with different individual amplitudes as well as relative phases to the mean-field (depicted in dashed black line).

We then analyzed what happens when oscillators with *different* phase space twist values are coupled through their mean-field. For this purpose, we took a system of 3 oscillators (with twist values *ε* = 0, *ε <* 0 and *ε >* 0) and turned on mean-field coupling after a certain time. We observed that all three oscillators synchronize to the mean-field for low values of the absolute value of twist (Figure 7B) although, due to the coupling-induced period differences, the relative phases of the individual oscillators to the mean-field depend on the specific twist values of the individual clocks: those with negative twist stay ‘behind’ the mean-field but those with positive twist oscillate with a phase advance (Figure 7B). Very large absolute values of twist, interestingly, produce again complex chaotic dynamics. The oscillator with a large value of *|ε|* does not synchronize to the mean-field and, as a result of the inter-oscillator coupling, this desynchronized oscillator ‘pulls’ to desynchronization the other two oscillators which otherwise would have remained in sync (Supplementary Figure S6).

## III. Discussion

This study aimed at characterizing period-amplitude correlations across circadian clock models of different complexity. Body clocks have to cope with cellular heterogeneity, what results in cellular clocks being variable across networks and tissues. We addressed whether the amplitude-period correlations that have been observed experimentally can be explained through heterogeneity in the cellular clocks, and what design principles are needed to produce such twist effects that we term *parametric twist*. Moreover, clocks live in constantly changing environments, that requires them to get adapted in the face of external changes. With the concept of *phase space twist* we study how this parameter, included explicitly in models, tunes the oscillator’s response to coupling, entrainment and pulse-like perturbations. We also retrieve the ‘old’ terminology of hard *versus* soft oscillators to refer to oscillations with negative and positive amplitude-period correlations, respectively.

It is important to note that a limit cycle oscillates with a characteristic period and amplitude regardless of the initial conditions. In contrast, conservative oscillations, like the Duffing oscillator (Figure 2), have a period and amplitude defined by the initial conditions. For this reason we have not termed *twist*, in the context of conservative oscillations, neither *parametric* nor *phase space*. We refer to *parametric twist* as that type of amplitude-period correlations that appears when studying an *ensemble* of limit cycle oscillators with differences in their intrinsic properties (i.e., biochemical parameters) and *phase space twist*, as that type of transient amplitude-period correlation that an *individual* oscillator experiences when encountering an external stimulus that results in an adaptation of its period and amplitude.

Parametric twist effects appear when analyzing populations of cells and require nonlinearities in oscillator models. The feedback loops needed to generate oscillations, which are commonly modeled with non-linear terms [41, 42, 50, 51], result in variations of parameters producing oscillations of different but correlated amplitudes and periods. We found that the type of parametric twist (positive or negative) depends on a number of factors: (i) on the biochemical parameter being affected, since different model parameters control the oscillation properties (amplitude and period) differently (Figures 3 and 5); (ii) on the region within the Hopf bubble where the clock’s parameter set is at (Figure 4); (iii) on the type of model and variable being measured: simple models with single negative feedback loops show the same parametric twist effects for all variables because the parameter of interest has the same effect on the amplitude of all variables. In complex models with synergies of loops, a change in one parameter might increase the amplitude of one variable but decrease the amplitude of a second variable (Figure 5 and Supplementary Table S1). This highlights the challenge of defining circadian amplitude (and twist effects) to find a metric for the whole oscillating system. Our findings explain why some experimental studies have found positive twist [26], others negative [14], whereas some other works found very little correlations [61].

We also addressed the concept of *phase space twist*, a parameter that we (and others [14,37,52]) introduced in a generic Poincaré oscillator. This twist parameter *ε* refers to period-amplitude correlations within a single oscillator, and it is a measure of the modulation of both instantaneous amplitude and period of the clock as the perturbed clock returns to the steady-state rhythm (i.e., limit cycle). Zeitgeber pulses, recurring zeitgeber inputs or coupling can all be regarded as ‘perturbations’, as any of these inputs modifies the natural clock’s limit cycle in phase space. Given this, it is not surprising that phase space twist, which characterizes the adaptation to a perturbation, has significant implications for phase response curves (PRCs), entrainment and coupling.

We start by discussing the role of twist in PRCs. The twist parameter affects the extent of the phase shift after a zeitgeber pulse (Figure 6D-F): increasing the twist parameter in absolute value results in alterations in the phase response curves and consequently in the resetting properties of oscillators. In particular, increasing *|ε|* increases the amplitude of the PRC (Figure 6J-L) until the PRC is converted from a type 1 PRC to a type 0 PRC [19, 62]. Consistently, amplitudes [17, 18, 20, 21, 63] and periods [18] of oscillators have also been shown to modulate their resetting properties. In these examples, clocks with short periods [18] and small amplitudes (in mathematical models [17, 18, 20] or in experiments with reduced coupling [20, 21] or mutations [63] that disrupt the normal rhythmicity) are easier to reset.

One can easily extrapolate these findings to responses to jet lag. If twist modulates the extent of a phase shift in response to a perturbation, it will also be critical in the adaptation to jet lag. This has indeed been found by Ananthasubramaniam et al. [23]. Interestingly, the authors found that the instantaneous amplitude effects induced by jet lag are also modulated by twist. In particular, positive twist aided recovery to jet lag in simple Goodwin-like models: reduced amplitudes were accompanied by faster clocks (i.e., shorter periods) upon a phase advance (e.g. when traveling eastwards), but larger amplitudes coincided with longer periods when the phase had to be delayed (e.g. when traveling westwards). Consistent with these theoretical observations are experiments performed in mice, where compromising coupling in the SCN (and thus decreasing amplitudes) reduce jet lag drastically, since resetting signals are much more efficient [21].

Transient amplitude-period correlations also affect the entrainment properties of an oscillator. Twist increases the range of zeitgeber periods to which the Poincaré oscillator can entrain (resulting in larger entrainment ranges, i.e., wider Arnold tongues, Figure 6M-O) and skews the resonance curves (Figure 6P-R). Some studies have even found coexisting limit cycles for driven oscillators with twist [32, 64]. Of note, however, is that the period of the Poincaré oscillator in all our phase space twist simulations was set to 24 h. Other theoretical studies have shown that clocks with different intrinsic periods show different amplitude responses to entrainment [23, 65], thus combining phase space and parametric twist effects.

Single cells harbor self-sustained clocks, but they are coupled (see [66] for a review on coupling) to produce a coherent rhythm at the level of tissues and organisms. Coupled networks show different rhythmic properties than clocks in isolation: phases synchronize and ensemble amplitudes increase for over-critical values of coupling strengths [59]. Our simulations provide an additional level of regulation, by showing that twist influences period length of individual clocks due to coupling. Positive twist results in periods lengthening upon mean-field coupling (Figure 7). The presence of coupling-induced changes in the oscillation period or amplitude can provide insights into the underlying oscillator type and the presence of twist. For instance, the longer period that has been observed in dispersed (and presumably uncoupled) U-2 OS cells compared to high-density cultures [67], but not in dispersed SCN neurons in culture [68, 69], could be explained by different implicit oscillator twist and amplitude relaxation rate values in different tissues.

In addition, the twist-induced modulation of period also affects the phase relation with which the clock oscillates respect to the mean-field: clocks with positive twist that tend to period-lengthen in response to coupling show later phases than those with negative twist, which tend to run quicker, consistent with chronotypes. It should be noted, however, that different phase relationships to the mean-field can also be obtained without twist, in networks of oscillators with different intrinsic periods [59]. But here again, the slower running clocks will tend to be phase-delayed in comparison to the faster-running clocks which will be phase-advanced. This suggests that twist is critical in how synchronization arises in network of coupled oscillators.

Our simulations assume that the coupling strength is constant across all oscillators. This assumption however might be questioned, since oscillators might ‘talk’ with different strengths to each other, especially because individual clocks are spatially organized within tissues, and have been shown to produce waves of oscillations for example in the SCN [70–72]. Thus, a more plausible scenario may involve local coupling, where clocks couple to neighbor clocks with a strength proportional to the oscillator distance. In fact, spatial gradient in nearest neighbor coupling can lead to a robust phase patterning [59]. Thus twist might not only be critical in how temporal synchronization but also spatial patterning arises in coupled ensembles.

The focus of this paper has been predominantly on twist, but it is important to remark that also the relaxation rate *λ* (i.e., how rigid/plastic an oscillator is) affects the response of oscillators to perturbations, consistent with previous computational work [20, 53]. It is in fact the ratio of *ε* to *λ* what determines the skewing of isochrones (see the analytical derivation in Materials and Methods) and the oscillator’s response to zeitgebers. Larger values of *λ* (more ‘rigid’ oscillators) imply that any perturbation is attracted back to the limit cycle at a higher rate and thus, perturbations ‘spend less time’ outside the limit cycle. Consequently, the twist parameter *ε* has a shorter effective time to act on the perturbed trajectory, and the isochrones become more straight and radial. Relaxation rate also has implications on coupling and entrainment. Rigid oscillators with *ε ≠* 0 display less coupling-induced amplitude expansions (Supplementary Figure S5B, [20]) and less period variations (Supplementary Figure S5A) in response to coupling than more ‘plastic’ oscillators with lower *λ* values. Moreover, rigid oscillators have smaller ranges of entrainment and narrower Arnold tongues [20]. Again, our results imply that any observation of resonant behavior and/or period changes might provide information on whether the underlying oscillator is a rigid *versus* plastic and hard *versus* soft clock.

We finish with an open question. Throughout our work we have claimed that period-amplitude correlations are widespread in both *in vivo* and in *in silico* clocks, and how they are critical to define how oscillators function in their environment. But, how does one integrate the circadian clock’s property of temperature compensation within this framework? The circadian clockwork is temperature-compensated [18, 44, 73–75], which means that increasing temperatures do not speed up significantly the clock, which still runs at approximately 24 h. A proposed hypothesis for temperature compensation suggests that temperature-sensitivity of circadian oscillation amplitude could stabilize the period [18, 75]. In support of this supposition is evidence from *Neurospora* [18] and *Gonyaulax* [76] suggesting that amplitude increases in response to temperature, but decreases in *Drosophila* [77]. What are the twist effects behind this adaptation? Temperature might affect how fast that variation is sensed in the clockwork and modulate the adaptation to find the new steady-state rhythm of larger or smaller amplitude at the new temperature.

## IV. Limitations and Conclusions

It is important to acknowledge that the amplitude-phase model proposed in this work exhibits radial symmetry and sinusoidal oscillations, but alternative models might not have these symmetry properties [53, 78]. It is worth noting that disparities in the underlying assumptions of models can significantly affect how an oscillator will interact with the environment, consequently affecting the oscillator’s response to zeitgeber pulses, coupling or entrainment.

Despite the complexities in quantifying amplitude, our models stress the important role of circadian amplitudes and their correlation with oscillator period. Although amplitudes are known to be regulated by a number of internal and external cellular factors, including cellular biochemical rates, light conditions [79], coupling [59], genetic and epigenetic factors [80] or ageing [81], our findings stress that twist effects (i.e. co-modulations of amplitudes and periods) also ‘feed back’ and affect the interaction of oscillators with the environment, facilitating entrainment, fastening response to pulse-like perturbations or modifying the response of a system to coupling. The theory of our conceptual models can also be applied to other oscillating system such as cardiac rhythms, somite formation, central pattern generators or voice production.

## V. Materials and Methods

### Conservative linear and non-linear oscillators: Harmonic vs. Duffing oscillators

In classical mechanics, a mass-spring harmonic oscillator is a system of mass *m* that, when displaced from its equilibrium position, experiences a restoring force *F* proportional to the displacement *x*, namely *F* = *−kx*, where *k* is the spring constant. When *F* is the only restoring force, the system undergoes harmonic motion (sinusoidal oscillations) around its equilibrium point. In the absence of damping terms, the harmonic motion can be mathematically described by the following linear second order ordinary differential equation (ODE)

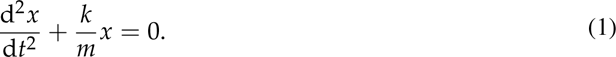

The solution to this differential equation is given by the function 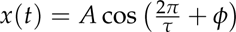 [30, 31], where *A* represents the amplitude; *φ*, the phase; and *τ* represents the period of the motion 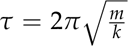. Thus the oscillatory period is determined only by the mass *m* and the spring constant *k*. The amplitude *A*, on the other hand, is determined solely by the starting conditions (by both initial displacement *x* and velocity 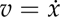).

Introducing a non-linear term in the restoring force such that *F* = *−kx − βx*^3^ allows the conversion of the simple harmonic oscillator into a Duffing oscillator [32]. Depending on the sign of *β*, the coefficient that determines the strength of the non-linear term, the spring is termed *hard* or *soft* oscillator. The equation of the Duffing oscillator, in the absence of damping terms, reads

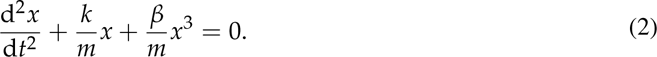

Due to the non-linear term introduced in equation 2, it is helpful to write this system in a form that can be easily treated via numerical integration. Considering the following change of variable 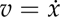, the equation can be reformulated as a system of two first order ODEs:

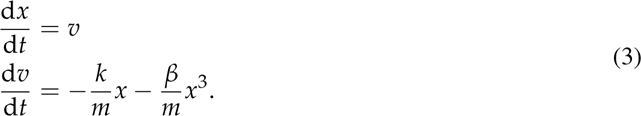

### Goodwin-like models

The three-variable Goodwin model is a minimal model based on a single negative feedback loop that can describe the emergence of oscillations in simple biochemical systems. All synthesis and degradation terms are linear, with the exception of the repression that *z* exerts on *x* which is modeled with a sigmoidal Hill curve. The equations that describe the dynamics read

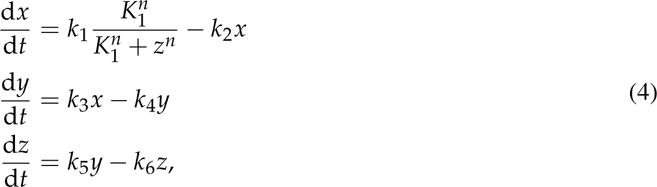

where *k*_1_, *k*_3_ and *k*_5_ represent the rates of synthesis of *x*, *y* and *z*, respectively; *k*_2_, *k*_4_ and *k*_6_, the degradation rates; and *n*, the Hill exponent.

The Goodwin model, however, requires a very large Hill exponent (*n* > 8) to produce self-sustained oscillations [48], which biologists and modelers have often considered unrealistic. Gonze [41] and others have shown that, by introducing additional nonlinearities in the system, the need for such high value of the Hill exponent can be reduced. In contrast to the linear degradation of variables of the original Goodwin model, the Gonze model [41] describes degradation processes with Michaelis-Menten kinetics as follows

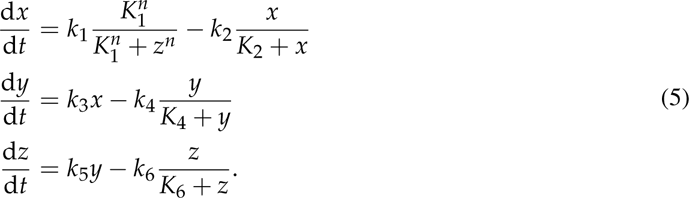

To mimic oscillator heterogeneity and evaluate the amplitude-period correlations among ensembles of clocks, degradation rates were varied around *±*10% their default value (Table 1).

### Almeida model

The Almeida model [51] is a protein model of the mammalian clockwork that includes 7 core clock proteins along with the PER:CRY complex and the regulation (activatory or repressive) that these exert on DNA binding sites known as clock-controlled elements to regulate circadian gene expression. The regulation at these clock-controlled elements, namely E-boxes, D-boxes and RORE elements, is described by the following terms:

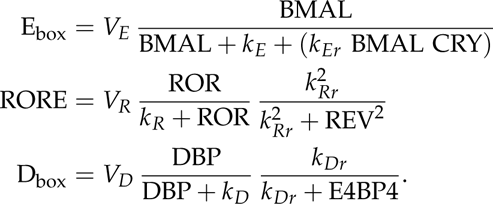

The system of ODEs that describes the dynamics of the clock proteins in the Almeida model reads

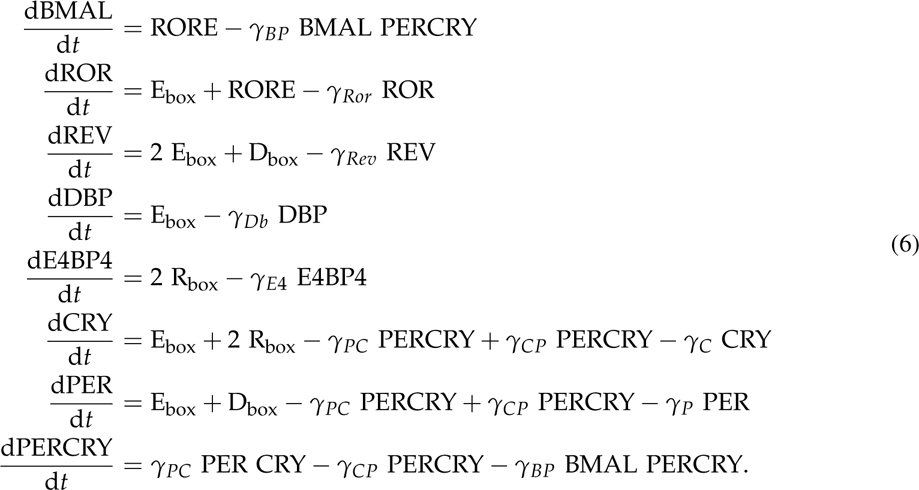

All parameter descriptions along with their default values are given in Table 2.

Parametric twist effects among ensembles of heterogeneous clocks were analyzed from simulations where all 18 parameters were randomly varied around *±*20% their default value (Table 2) one at a time. Since this model can lead to period-doubling effects upon changes of certain parameters [82], those oscillations whose period change resulted in oscillations with period-doubling were removed from the analysis. Moreover, if changing the default parameter in the ensemble resulted in a ratio of amplitude variation relative to the default amplitude whose range was < 0.1, then we considered that ensemble to have no twist for that particular control parameter (Supplementary Table S1).

### Poincaré model

The intrinsic dynamical properties from single oscillators and their interaction to external stimuli can be very conveniently described by means of a Poincaré model [37]. We here propose a modification of its generic formulation that explicitly takes into account twist effects through the phase space twist parameter *ε*. The modified Poincaré model with twist reads

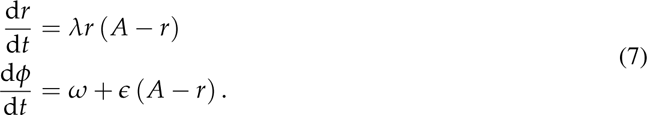

The first equation describes the rate of change of the radial coordinate *r*(*t*) (i.e., the time-dependent distance from the origin), whereas the second equation determines the rate of change of the angular coordinate *φ*(*t*), where 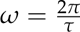. The parameters *τ*, *A*, *λ* and *ε* denote the free-running period (in units of time), amplitude (arbitrary units), amplitude relaxation rate (in units of time^-1^) and phase space twist of the oscillator (in units of time^-1^), respectively. In the absence of twist, namely *ε* = 0, the phase changes constantly along the limit cycle at a rate 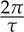, independently of the radius. In the case of *ε* ≠ 0, the phase changes at a constant rate only when *r* = *A*; if any perturbation is to modify *A* such that *r ≠ A*, then the phase change will be accelerated or decelerated depending on the sign of *ε* and on whether *r > A* or *r < A*. The model parameters, unless otherwise specified in the figures or captions, are the following: *A* = 1, *λ* = 0.05 h^-1^, *τ* = 24 h and *ε* values of 0 or *±*0.1 h^-1^.

To study the effects of twist on the oscillator’s response to pulse-like perturbations pert(*t*), periodic zeitgeber input *Z*(*t*) or mean-field coupling *M*, the individual Poincaré oscillators *i* were converted into Cartesian coordinates and the respective terms were added in the equations of the *x_i_*variable as follows:

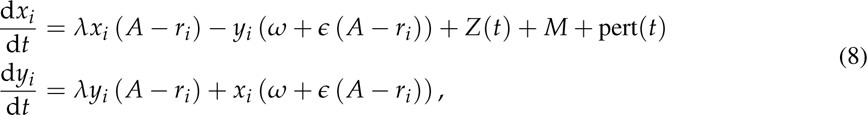

where 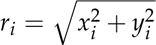. The zeitgeber *Z*(*t*) is given by:

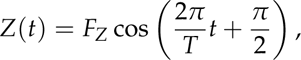

where *T* represents the zeitgeber period and *F_Z_* the strength of the zeitgeber input. The mean-field *M* coupling is given by:

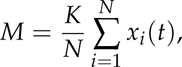

where *K* represents the coupling strength and *N* is the number of oscillators in the coupled ensemble. Lastly, the square-like perturbation pert(*t*) is defined as follows:

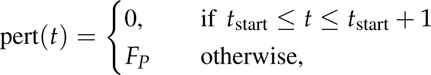

where *F_P_* is the strength of the perturbation and *t*_start_ is the time at which the perturbation starts. The perturbation lasts 1 h and is set to *F_P_* = 0.7 a.u. in all our simulations.

#### Analytical derivation of isochrones

Arthur T. Winfree introduced the concept of isochrones as any set of dynamical states which oscillate with the same phase when they reach the limit cycle at time *t →* ∞ [52]. That asymptotic phase at *t →* ∞ is what Winfree termed latent phase Φ. Since the dynamical flow of the Poincaré model has polar symmetry, the isochrones must also have polar symmetry such that

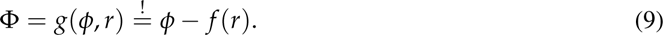

Moreover, the latent phase velocity Φ oscillator follows its kinetic equation: necessarily increases at units of the angular velocity 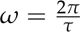 as the oscillator follows its kinetic equation:

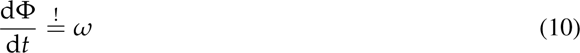

Combining the two previous equations and with the use of the chain rule, we can calculate the radius dependency of the latent phase Φ:

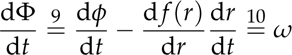

The terms 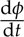 and 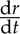 are defined in the Poincaré model (equation 7), so that the equation above can be rewritten as:

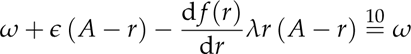

Solving for 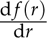:

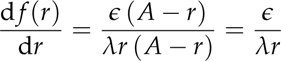

Next, the solution for *f* (*r*) can be found through integration:

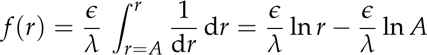

If *A* = 1, the last term can be neglected such that

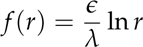

Finally, the solution of *f* (*r*) can be inserted in equation 9 to end up with the equation for isochrones as a function of the radius that we have used to simulate Figure 6:

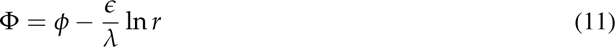

Isochrones are thus loci of polar coordinates (*φ*, *r*) in phase space with the same latent phase Φ. Equation 11 shows how the shape of the isochrone for the Poincaré model in equation 7 depends on the ratio of twist *ε* to relaxation rate *λ*. Specifically, higher *ε* values (in modulus) and lower relaxation rates *λ* lead to more curved isochrones (Supplementary Figure S1).

### Computer simulations and data analysis

All numerical simulations were performed and analyzed in Python with the `numpy`, `scipy`, `pandas` and `astropy` libraries. The function `odeint` from `scipy` was used to numerically solve all ordinary differential equations. Bifurcation analyses for Figure 3C, D, Figure 4 and Figure 5B, E were performed in XPP-AUTO [83] using the parameters *N_tst_* = 150, *N_max_* = 20000, *Ds_min_* = 0.0001 and *Ds_max_* = 0.0002.

Throughout our analyses, periods were determined by (i) normalizing the solutions to their mean, (ii) centering them around 0 (by subtracting one unit from the normalized solution), and (iii) by then computing the zeroes of the normalized rhythms. The period was defined as the distance between two consecutive zeros with a negative slope. Amplitudes were determined as the average peak-to-trough distance of the last oscillations after removing transients.

## Abbreviations

SCN: suprachiasmatic nucleus
PRC: phase response curve
ODE: ordinary differential equation.

## Acknowledgments

The authors thank Bharath Ananthasubramaniam, Gianmarco Ducci, Jihwan Myung and Adrian Granada for stimulating and helpful comments as well as constructive criticism. The authors would also like to acknowledge the work done by the Bachelor and Master students Pia Rose, Julia K. Schlichting, Barbara Zita Peters Couto and Rebekka Trenkle.

## Data accessibility

The source code used to generate the simulated data and reproduce all figures is available through GitHub (https://github.com/olmom/twist).

## Author contributions

Study design and conceptualization: M.dO. and H.H.; Methodology: M.dO., C.S., and H.H.; Investigation: M.dO., C.S. and H.H.; Resources and data curation: M.dO., C.S., C.M., and H.H.; Writing (original draft): M.dO. and H.H.; Writing (review & editing): M.dO., C.S., S.G., C.G., A.K. and H.H.; Funding acquisition: A.K. and H.H.

