## Supplementary Material: Table S1 and Figures S1-S6 for "Are circadian amplitudes and periods correlated? A new *twist* in the story"

**Table S1: Summary of parametric twist effects upon changes of model parameters in the Almeida model.** Model parameters were changed individually around ±20% their default parameter value (see Table 2 in main text) to simulate oscillator heterogeneity and study amplitude-period correlations from all variables of the ensemble. Due to the synergies of feedback loops present in the Almeida model [1], different parametric twist effects can appear for variations of one particular parameter depending on the measured variable. If the default parameter variation resulted in an amplitude-period correlation where the range of ratio of amplitude variation (compared to the default amplitude) was $< 0.1$, that ensemble was considered to have no twist (twist = 0) for that particular control parameter. + and – signs refer to the sign of the correlations.

| | **±10% parameter change in** | | | | | | | | | | | |
|---|---|---|---|---|---|---|---|---|---|---|---|---|
| effect on: | $V_R$ | $k_R$ | $k_{Rr}$ | $V_E$ | $V_D$ | $\gamma_{Ror}$ | $\gamma_{Rev}$ | $\gamma_P$ | $\gamma_C$ | $\gamma_{PC}$ | $\gamma_{CP}$ | $\gamma_{BP}$ |
| **BMAL1** | + | + | + | – | – | 0 | 0 | + | + | + | 0 | 0 |
| **ROR** | 0 | 0 | – | 0 | 0 | 0 | – | 0 | 0 | 0 | 0 | 0 |
| **REV** | + | + | 0 | + | 0 | 0 | + | + | + | 0 | 0 | 0 |
| **DBP** | + | + | + | 0 | 0 | + | + | + | 0 | + | 0 | 0 |
| **E4BP4** | 0 | 0 | 0 | 0 | + | + | 0 | 0 | 0 | 0 | 0 | 0 |
| **CRY** | 0 | 0 | 0 | 0 | 0 | 0 | 0 | 0 | 0 | 0 | 0 | 0 |
| **PER** | + | + | + | + | 0 | + | + | + | + | + | 0 | 0 |
| **PERCRY** | + | + | + | + | 0 | + | + | + | + | + | 0 | 0 |

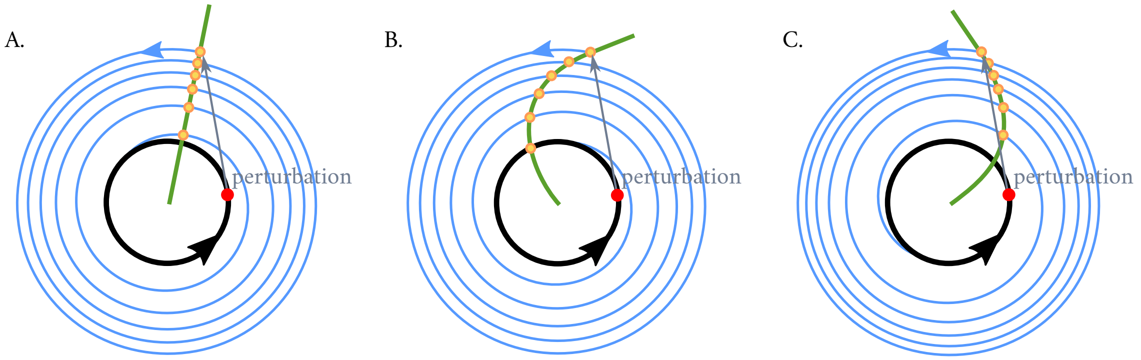


**Figure S1: Schematic of an isochrone in a Poincaré oscillator with (A) no twist, (B) positive twist or (C) negative twist.** To understand the concept of isochrones, a simple experiment is performed: if one considers a point in phase space (shown in red) within a limit cycle (black) and observes where the system returns to after exactly one period, the answer is trivial: to the same spot. If, however, now a perturbation is applied (grey arrow) such that the system starts in a section of the state space which is outside of the stable periodic orbit, and one maps the points that the system 'leaves as a footprint' after exactly one period as it relaxes back to the limit cycle (blue trajectories), these points (shown in yellow) will outline an isochrone (green). The twist parameter $\epsilon$ affects the curvature of the isochrone, with implications in the response of the oscillating system to the perturbation. The oscillator with positive twist (B) arrives at an earlier phase than that with no twist (A), resulting in a phase advance with respect the clock with no twist, whereas the clock with negative twist (C) arrives to the limit cycle at a later phase (i.e., delayed with respect to the clock with no twist).

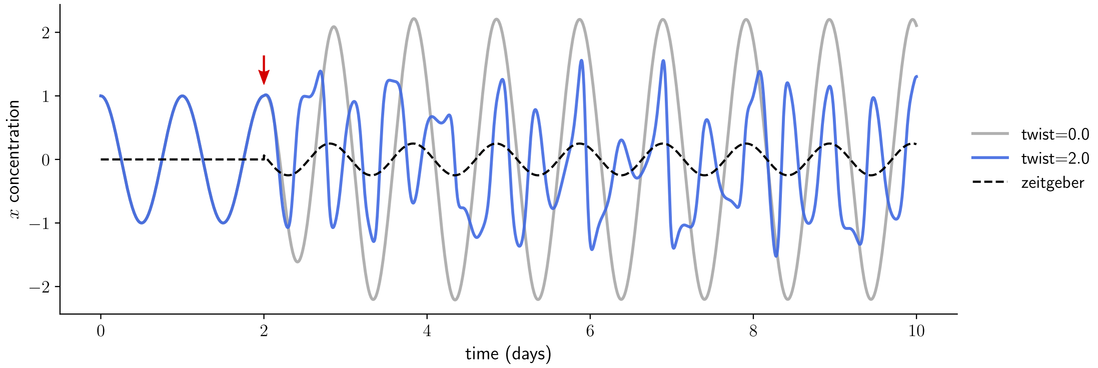


**Figure S2: Large values of oscillator twist $\epsilon$ affect sync of individual oscillators to external zeitgeber inputs.** An oscillator with no twist (shown in grey) entrains to the zeitgeber (indicated with a dashed line) when the periodic input is turned on (red arrow, $T = 24.5$ h, $F = 0.25$), but the oscillator with large positive twist (blue) does not entrain and loses its periodicity. Both oscillators, besides their twist value, are otherwise identical (same free running period $\tau = 24$ h, amplitude $A = 1$ and amplitude relaxation rate $\lambda = 0.05\,\mathrm{h}^{-1}$).

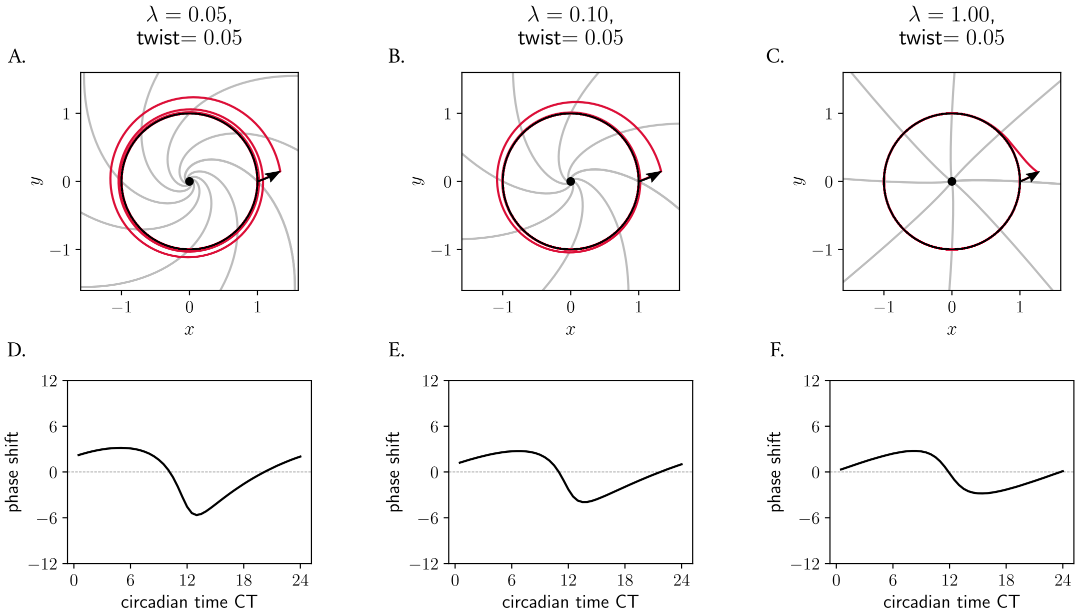


**Figure S3: Role of amplitude relaxation rate on the skewing of isochrones and consequently on the response of oscillators to perturbations**. (A-C) Poincaré oscillator models with different amplitude relaxation values $\lambda$, but otherwise identical (free running period $\tau = 24\,\mathrm{h}$, amplitude $A = 1$ and twist $\epsilon = 0.05\,\mathrm{h}^{-1}$), shown in phase space. The limit cycle is shown in black; perturbed trajectories are shown in red; isochrones are depicted in grey. (D-F) Phase response curves: the shape of the PRC and the extent of the phase shifts depends on the relaxation rate $\lambda$. See Materials and Methods for details on the analytical derivation of the equation of isochrones and roles of $\lambda$ and $\epsilon$.

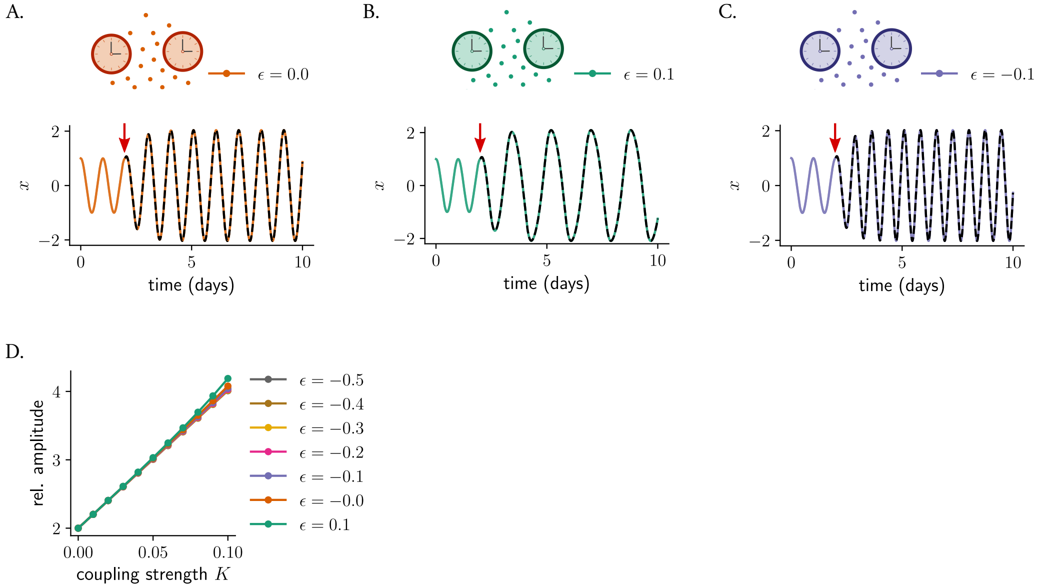


**Figure S4: Coupling induces amplitude expansions and period changes that depend on the twist $\epsilon$.** Effect of mean-field coupling on two oscillators with (A) $\epsilon = 0$, (B) $\epsilon = 0.1$ or (C) $\epsilon = -0.1$ but otherwise identical (free running period $\tau = 24\,\text{h}$, amplitude $A = 1$ and amplitude relaxation rate $\lambda = 0.05\,\text{h}^{-1}$). The time series show how the period of the coupled network depends on the twist parameter $\epsilon$. The moment in which mean-field coupling is turned on, at a coupling strength $K = 0.1$, is shown with a red arrow; the mean-field is shown with a dashed black line. (D) The amplitude of the coupled network increases with coupling strength $K$, but with no major differences for networks with different twist values.

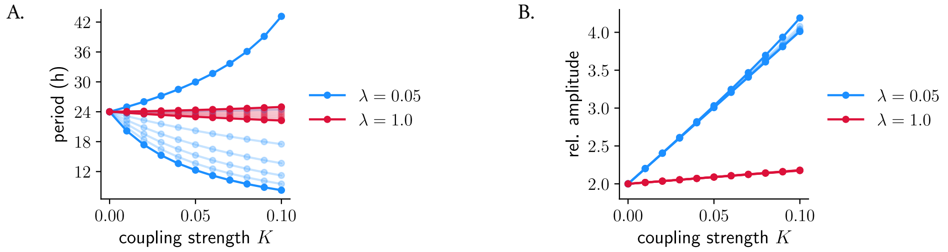


**Figure S5: Role of amplitude relaxation rate on coupling-induced period and amplitude changes.** More rigid clocks (higher $\lambda$ values, red lines) are less sensitive to twist-induced effects and thus display less coupling-induced period changes (A) or amplitude expansions (B) than weaker clocks (in blue).

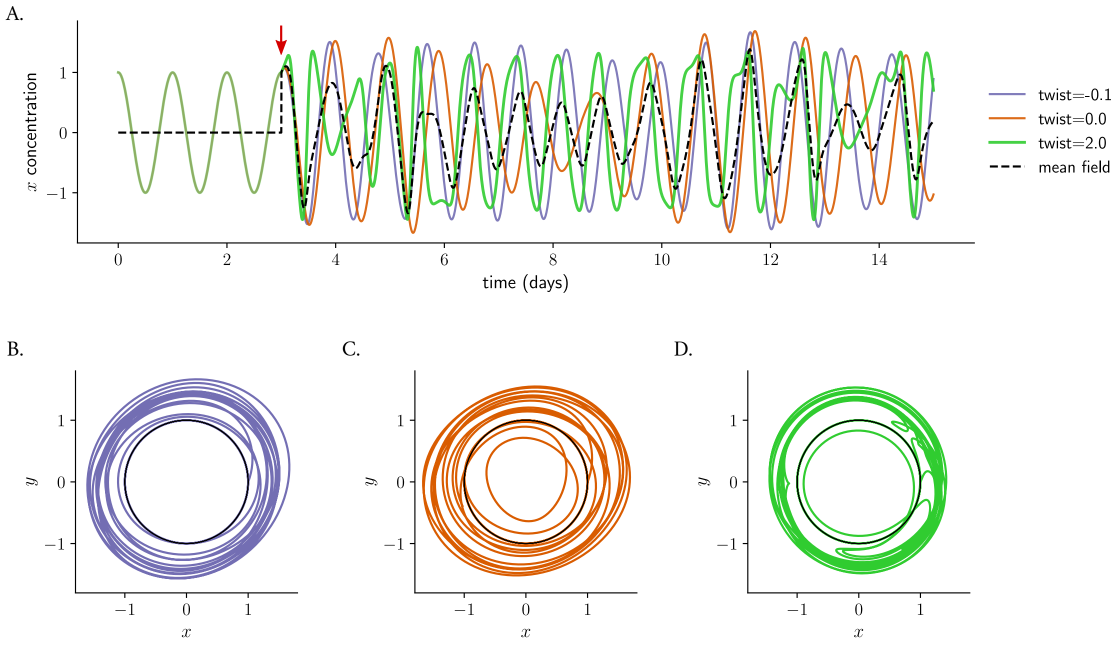


**Figure S6: Twist-induced chaos upon mean-field coupling.** (A) Oscillators with $\epsilon = -0.1, 0, 2$ oscillate in a self-sustained manner in the absence of coupling, but when mean-field coupling at $K = 0.1$ is turned on (red arrow), periodicity is lost. The oscillator with $\epsilon = 2$ (green line) cannot synchronize due to its large twist and thus affects the rest of the network (purple and orange lines become arrhythmic too). As a result, the mean-field (shown in black and with dashed line) also becomes arrhythmic. (B-D) Twist-induced chaotic trajectories of the oscillators from (A) shown in phase space. The limit cycles from the uncoupled clocks are shown in black.
